# Antibiotic persistence does not cause phenotypic heterogeneity in tolerance of *Escherichia coli* to formaldehyde stress but can preserve it through time

**DOI:** 10.1101/2022.09.23.509177

**Authors:** Isaiah D. Jordan, Tomislav Ticak, Jessica A. Lee, Christopher J. Marx

**Author notes:** Corresponding author: Department of Biological Sciences, University of Idaho, Moscow, ID, USA.

## Abstract

The phenomenon of phenotypic heterogeneity, where an isogenic population expresses varying phenotypes, has been uncovered for an expanding variety of organisms and traits. This heterogeneity in phenotype can be physiologically relevant, such as the ability for antibiotic persistence to allow populations to survive lethal conditions due to rare cells being in a slow or non-growing state. Recently, it was discovered that *Methylorubrum extorquens*, a facultative methylotroph, possesses a continuous spectrum of phenotypic states conferring tolerance to formaldehyde. Formaldehyde is a toxin produced by *M. extorquens* during growth on methanol. The phenotypic tolerance allowed rapid growth on levels of formaldehyde which were lethal to the majority of the population. Transcriptomics indicated this may be due to upregulation of proteins that attenuate oxidative stress and protein damage rather than increasing formaldehyde oxidation to prevent accumulation. These data suggested that heterogeneity to formaldehyde stress may be present even in non-methylotrophic organisms that do not routinely produce large quantities of formaldehyde, and thus widely distributed across bacteria. To investigate this, we tested *Escherichia coli* for heterogeneity to formaldehyde stress during growth on glucose. Like *M. extorquens, E. coli* populations have a wide, continuous range of formaldehyde tolerance thresholds and this tolerance was reversible. Several other features, however, were different from what was found for *M. extorquens*. Most *E. coli* growth occurred after formaldehyde levels had dropped, suggesting that persistence could be the cause. The dynamics of antibiotic persistence and formaldehyde tolerance were both tracked but found to not be correlated. On the other hand, the persister cell state can maintain formaldehyde tolerance. These data suggest that persistence can preserve phenotypic heterogeneities in other traits, further expanding its potential role in helping cells survive environmental stressors.

## Introduction

Isogenic bacterial cultures are often assumed to be comprised of individuals that are identical and interchangeable in phenotype, but the validity of this assumption is increasingly being challenged. It is now recognized that there can be substantial “phenotypic heterogeneity” whereby single cells of genotypically identical organisms express different phenotypes. Indeed, far from being a rare, incidental phenomenon, phenotypic heterogeneity has been observed in many species [1], and within a species it is not uncommon for many traits to be heterogeneous [2] [3]. It seems likely that with increasing awareness of phenotypic heterogeneity and interest in its implications the list of species and traits that it affects will continue to grow.

Phenotypic heterogeneity can form distributions that are either discrete or continuous. Perhaps the most familiar scenario to microbiologists is the case in which there are discrete cell forms, such as flagellated and non-flagellated bacteria [4], stalked and swarmer cells of *Caulobacter* [5], or nitrogen fixing heterocysts interspersed along photosynthetic cyanobacterial filaments [6]. More recently, approaches such as transcriptional reporters have revealed continuously heterogeneous traits that lie across a wide spectrum of trait values. A striking example of this was seen in a study by Elowitz and colleagues, where two fluorescent genes were expressed in *Escherichia coli* under a pair of identical, constitutive promoters, resulting in individuals representing all combinations of the two colors existing in the culture [7]. Because these expression ratios existed within single cells, this allowed the authors to rule out an extrinsic factor between cells as the source of variation, such as spatial heterogeneity in the environments. This revealed that the observed expression heterogeneity resulted from intrinsic factors. There are many potential sources of intrinsic factors, such as stochasticity in gene expression [7],[8], asymmetric inheritance of the proteome [9], or distinct DNA methylation states [10], [11].

Phenotypic heterogeneity generates a fuzzy mapping of genotypes to phenotype, which under some conditions can provide a fitness advantage. A well-characterized example where phenotypic heterogeneity results in survival of lethal stressors is antibiotic persistence. Whereas antibiotic resistance represents genetic changes that allow all cells to survive a given dose, persistence is caused by purely phenotypic differences between cells. As first noted in 1944 [12], killing with certain antibiotics, such as penicillin or ampicillin, first leads to a rapid exponential decline in viable cell counts, only to be followed by a period of much slower killing. These persistent cells are now known to have entered a non-growing (or very slow growing) state prior to exposure to the stressor, rendering them unable to increase in number, but able to avoid or reduce susceptibility to a wide variety of antibiotics and other toxic insults to the cell [13]. Once the concentration of the antibiotic declines, these cells can safely and spontaneously transition back into a growing state and re-establish the population. During exponential growth, the proportion of persisters tends to be quite low (~10^-3^), but these numbers increase dramatically upon nutrient depletion and entry into stationary phase [14], [15]. Critically, the distribution of growth rates between growing cells and persisters, as well as in their phenotypic tolerance to toxins, are each bimodal (i.e., not a wide spectrum of growth rates and tolerances) and are perfectly anticorrelated. As suggested by the above-mentioned bimodal distribution of growth rates, persistence consists of two distinct, discrete states which cells can transition between [16].

Recently, our laboratory discovered a novel phenomenon of phenotypic heterogeneity in populations of *Methylobacterium extorquens* [17]. *M. extorquens* PA1 is a facultative methylotrophic alphaproteobacterium which, during growth on methanol, oxidizes it to formaldehyde as the first metabolic intermediate [18],[19], [20]. This formaldehyde is then oxidized to formate [21], [22], which is then either dissimilated to CO_2_ or assimilated into biomass [23], [24]. This work found that the ability of cells to grow on methanol in the presence of additional external formaldehyde was quite variable within the population. Rather than two, discrete subpopulations, there was a wide spectrum of maximal formaldehyde concentrations that individual cells could tolerate and still grow at typical rates. In some ways this phenomenon was similar to persistence: no genotypic differences, tolerance was pre-existing prior to the addition of stressor, and the phenotype was reversible. But in other important ways it is quite distinct, in that tolerant cells survive stress while actively growing in the presence of otherwise lethal levels of formaldehyde and they exist along a continuum of tolerance levels.

As a first step towards understanding the physiological basis of phenotypic heterogeneity in formaldehyde tolerance, the transcriptome of populations with high and low tolerances were sequenced (17). Surprisingly, transcription of formaldehyde oxidation genes was unchanged in tolerant cells. Rather, upregulated genes were mostly related to the management of oxidative stress and protein damage, indicating that formaldehyde tolerance in *M. extorquens* may be less a result of heterogeneity in the ability to degrade the toxin and more dependent upon differences between cells in managing its negative effects.

Because phenotypic heterogeneity for formaldehyde tolerance in *M. extorquens* was apparently due to general, well-conserved cellular mechanisms, we hypothesized that this heterogeneity may also manifest in other species of bacteria, such as *E. coli*, which may also exhibit cell-to-cell differences in these functions. Here we describe the heterogeneity of tolerance to formaldehyde in *E. coli*, which both shares commonalities as well as differences from that seen for *M. extorquens*, namely in that growth does not occur until formaldehyde is degraded to very low levels. Additionally, we determined that antibiotic persistence is not responsible, despite the observed extended lag phase. However, we did find indications that persistence does play a role in preserving a heterogeneous phenotype of formaldehyde tolerance long after its induction had ceased, suggesting a role for persistence in preserving phenotypic heterogeneity in other traits.

## Methods and Materials

### Media and chemicals

Lysogeny broth (LB) (Sigma-Aldrich, St. Louis, MO) was used for pre-growth in liquid from single colonies on LB agar plates. Cultures were then grown overnight in a modified formulation of the minimal MOPS media [25] (see S1 Table) with 11.1 mM glucose.

### Strain and growth

*E. coli* strain NCM3722, a strain derived from the K-12 strain MG1655 [26] was used for all experiments. The stock was obtained from Terrence Hwa at the University of California San Diego and stored in 25% glycerol at −80 °C. All cells were incubated at 37 °C, whether in liquid or on plates. Fresh streak plates were created weekly by stabbing a pipette tip into the frozen stock and placing the tip in 5 mL lysogeny broth in a culture tube at room temperature until all ice on the tip had melted into the broth. The tube was then incubated for roughly six hours and then streaked for isolated colonies following overnight growth. Plates were stored at 4 °C with a parafilm seal for up to one week.

Liquid cultures were incubated in 5 mL of culture in a New Brunswick Innova 44 incubator at 250 rpm in capped 28 mL 18 × 150 mm culture tubes held in racks slanted at 45°. Most plates were incubated inverted until colonies were sufficiently large to be easily countable. In the case of formaldehyde treated plates, this necessitated checking and counting colonies multiple times due to highly heterogeneous colony arisal times. Plates grown on scanner beds were not inverted to allow for their imaging. Black felt was placed in the lids of these plates to both improve contrast for imaging and to absorb condensation to prevent drops of water falling on the agar. For ampicillin-treated cells, 200 μL of culture was incubated in Eppendorf microcentrifuge tubes with 100 μg/mL ampicillin for 2.5 hours (see S2 figure).

### Manual optical density readings

Optical density at 600 nm (OD_600_) was measured on a Bio-Rad SmartSpec Plus spectrophotometer and diluted with MOPS medium when necessary to stay within the linear range.

### Spot plating

For all experiments involving spot plating to determine colony forming units per mL, a series of 1/10 serial dilutions were performed in MOPS media without carbon in a 96 well plate. Three sets of 10 μL spots were then plated for each dilution using a multichannel pipettor on MOPS agar with glucose (or formaldehyde, if appropriate). For each set of spots, the two adjacent dilutions yielding a number of colonies that could be accurately determined were counted. They were then summed and divided by 11 and multiplied by the dilution factor of the more concentrated dilution. The counts from each set of spots were then averaged to estimate the concentration of colony forming units per mL in the culture.

### Growth analysis in multiwell plates

A BioTek Synergy H1 plate reader was used for multiwell growth analyses. 600 μL of inoculated media was dispensed into each well of a Corning Costar 48 flat bottom clear well plate. The plate was then shaken with a double orbital pattern with a 3 mm throw at 37 °C and 425 cpm, with readings automatically taken every 15 minutes. Breathe-Easy sealing membrane (Sigma-Aldrich, St. Louis, MO) was used to prevent evaporation, and a 1 °C gradient across the bottom and top of the plate was used to prevent condensation.

### Formaldehyde preparation and quantitation via Nash assay

One molar formaldehyde was prepared by mixing 0.3 g of paraformaldehyde (Sigma-Aldrich, St. Louis, MO) with 9.76 mL of ultrapure water (18.2 mΩ·cm), with 50 μL of 10 N NaOH (Sigma-Aldrich, St. Louis, MO), for a final volume of 10 mL. It was then boiled in a stoppered and crimp sealed Balch tube for 20 minutes and used for up to a week; fresh formaldehyde stock was prepared weekly. A needle and syringe were used to aseptically extract formaldehyde from the tube for use.

Formaldehyde concentrations were assessed using the Nash assay [27]. The assay is performed by mixing 250 μL of reagent B, which is 2 M ammonium acetate (Sigma-Aldrich, St. Louis, MO), 0.05 M acetic acid (Macron Fine Chemicals, Radnor, PA), and 0.02 M acetylacetone (Sigma-Aldrich, St. Louis, MO) in deionized, distilled water, with 250 μL of sample (potentially diluted as needed to stay within the linear range of the assay), vortexed or inverted briefly and incubated at 60 °C for 6 minutes. Absorbance at 412 nm was then measured on a Bio-Rad SmartSpec Plus spectrophotometer (Bio-Rad, Hercules, CA) immediately. For inoculated media, to avoid the possibility that cells would absorb at those wavelengths, 300 μL of media was spun for one minute at 20,817 *g* to pellet cells and then 250 μL of the supernatant used for the assay. Standard curves were generated using serial dilutions of 1 M formaldehyde stock prepared as described above.

### Formaldehyde oxidation enzyme assay

To measure differences in formaldehyde oxidation rates between lysates of cells grown under different conditions, cells were first lysed using B-PER Complete Bacterial Protein Extraction Reagent (Thermo Fisher Scientific, Waltham, MA). To do this, cultures were allowed to grow to mid log (OD_600_ ~ 1) and pelleted by centrifugation at 20,817 *g* for 10 minutes in 15 mL centrifuge tubes. The supernatant was poured off, and the pellets were stored at −80 °C until the assay was performed. Then, the pellet was pipetted into a microcentrifuge tubule and weighed. Five mL of the B-PER reagent was added for each gram of pellet, and 2 μL of 0.5 M EDTA (Sigma-Aldrich, St. Louis, MO) added per mL of B-PER. The mixture was then gently rocked for 15 minutes at room temperature. Following this, the microcentrifuge tube containing the mixture was centrifuged at 20,817 *g* for 20 minutes to spin down large remnants of cells. A sample of the supernatant was used to initiate a reaction in Bradford reagent (Thermo-Fisher Scientific, Waltham, MA) by adding 5 μL of sample to 600 μL of Bradford reagent in a one cm cuvette, inverting a few times and allowing the mixture to incubate for 10 minutes. Following incubation, the absorbance of the solution was measured at 595 nm with a 0.5 cm pathlength. A standard curve was created using the above protocol with Quick-Start Bovine Serum Albumin Standard (Bio-Rad, Hercules, CA) between 0.125 – 2 mg/mL to permit the quantification of protein in the sample. Samples outside the standard curve were rerun with dilutions to stay within the linear range of the assay.

Having determined the protein content of the sample, the capacity for formaldehyde oxidation via glutathione in each treatment was measured across a series of protein and formaldehyde concentrations. The reaction mixture in all wells consisted of 100 mM NaCl (VWR, Radnor, PA), 50 mM K_2_PO_4_ (VWR, Radnor, PA), 1 mM NAD^+^ (Roche Diagnostics, Basel, Switzerland), and 2 mM reduced glutathione (Sigma-Aldritch, St. Louis, MO). Additionally, for each treatment, 25, 12.5, 6.25 and 0 μg of protein were tested against 0.1, 0.05, 0.025 and 0 mM formaldehyde, for a total of 16 wells per treatment. The absorbance of each well with reaction mixture and protein (but without formaldehyde) was measured at 340 nm with a BioTek Synergy H1 plate reader (BioTek, Winooski, VT) in a Corning CoStar 48 well plate (Corning, Corning, NY). Following this, the reaction was initiated with formaldehyde, and shaken for 5 minutes before being read again. This step was repeated until a total of 9 cycles (45 minutes) had elapsed.

## Results

### Individuals have heterogeneous tolerance to formaldehyde stress

To investigate whether *E. coli* exhibits phenotypic heterogeneity in response to formaldehyde stress, cultures that had undergone various treatments were spot plated onto various concentrations of minimal glucose plates containing formaldehyde. In all treatments, increasing the amount of formaldehyde on the agar plates decreased the CFU in a continuous manner (Fig 1, orange line), suggesting that a wide spectrum of tolerances to formaldehyde stress exist in *E. coli*, with a small drop in viability by 0.4 mM formaldehyde, and further reductions at higher concentrations. The culture grown in media containing 0.8 mM formaldehyde had higher viability across this range and even had a small subpopulation of cells that could form colonies on agar with 1 mM formaldehyde (Fig. 1, pink line). Inoculum from this culture was regrown to stationary in media without formaldehyde and retested, revealing that its tolerance distribution transitioned back to resemble the naive population (Fig. 1, green line). This return to the original phenotype distribution, as was seen with *M. extorquens* [17], suggests that these are phenotypic and not genotypic variants. Furthermore, these data support that there is a continuous, phenotypically heterogeneous and plastic response to formaldehyde stress in E. coli.

**Figure 1.**
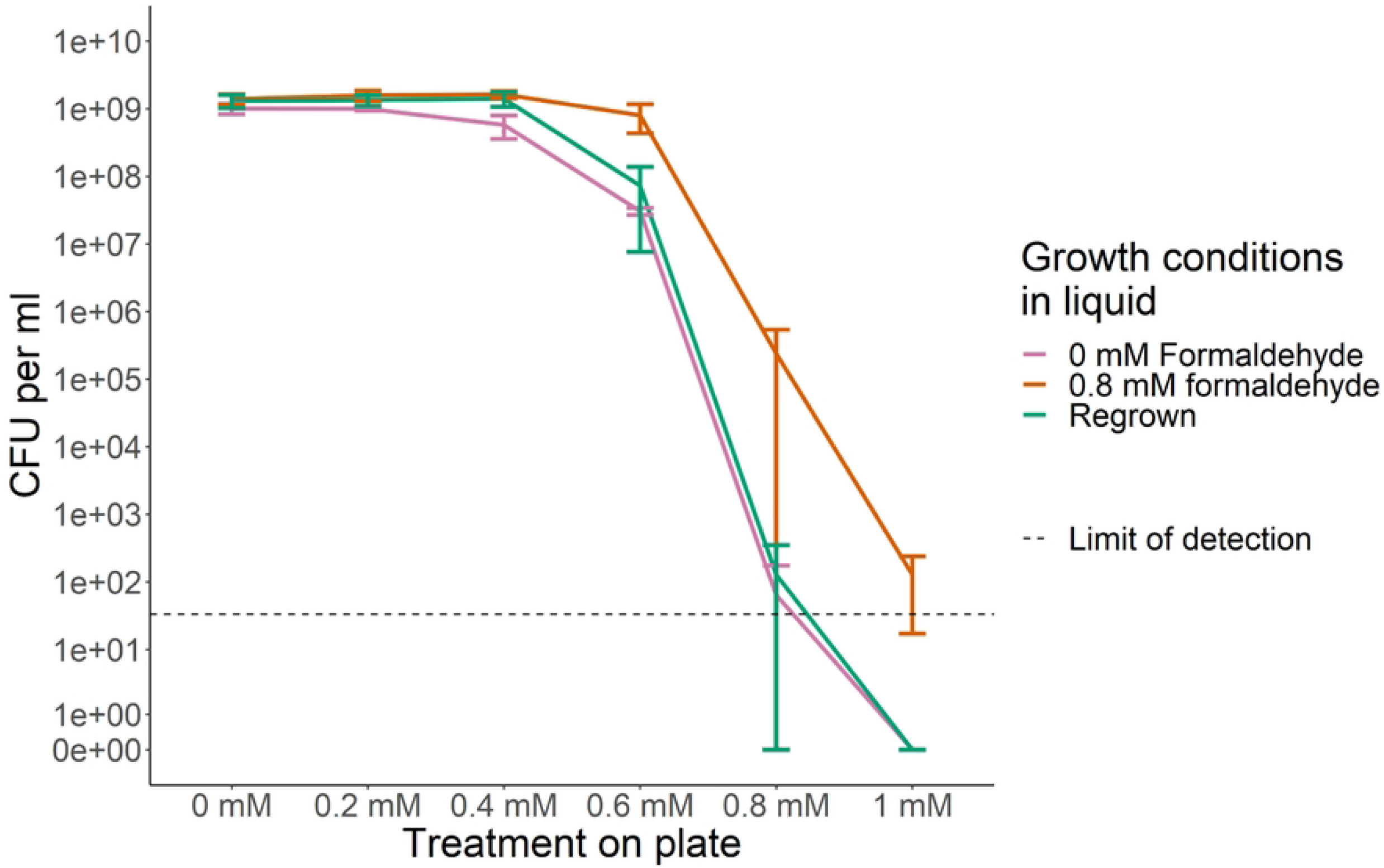
Rare tolerant cells can survive otherwise inhibitory concentrations of formaldehyde. An overnight culture of cells plated onto minimal glucose plates with various concentrations of formaldehyde exhibited a range of survival, suggesting some cells had enhanced tolerance to formaldehyde (pink). Cells pre-grown to stationary phase in medium with formaldehyde exhibited a shifted distribution of heterogeneity in tolerance (orange). Upon regrowth in the absence of formaldehyde, tolerance shifted back to its original distribution (green). The limit of detection (one colony observed) is shown by the dashed line.

**Figure 2.**
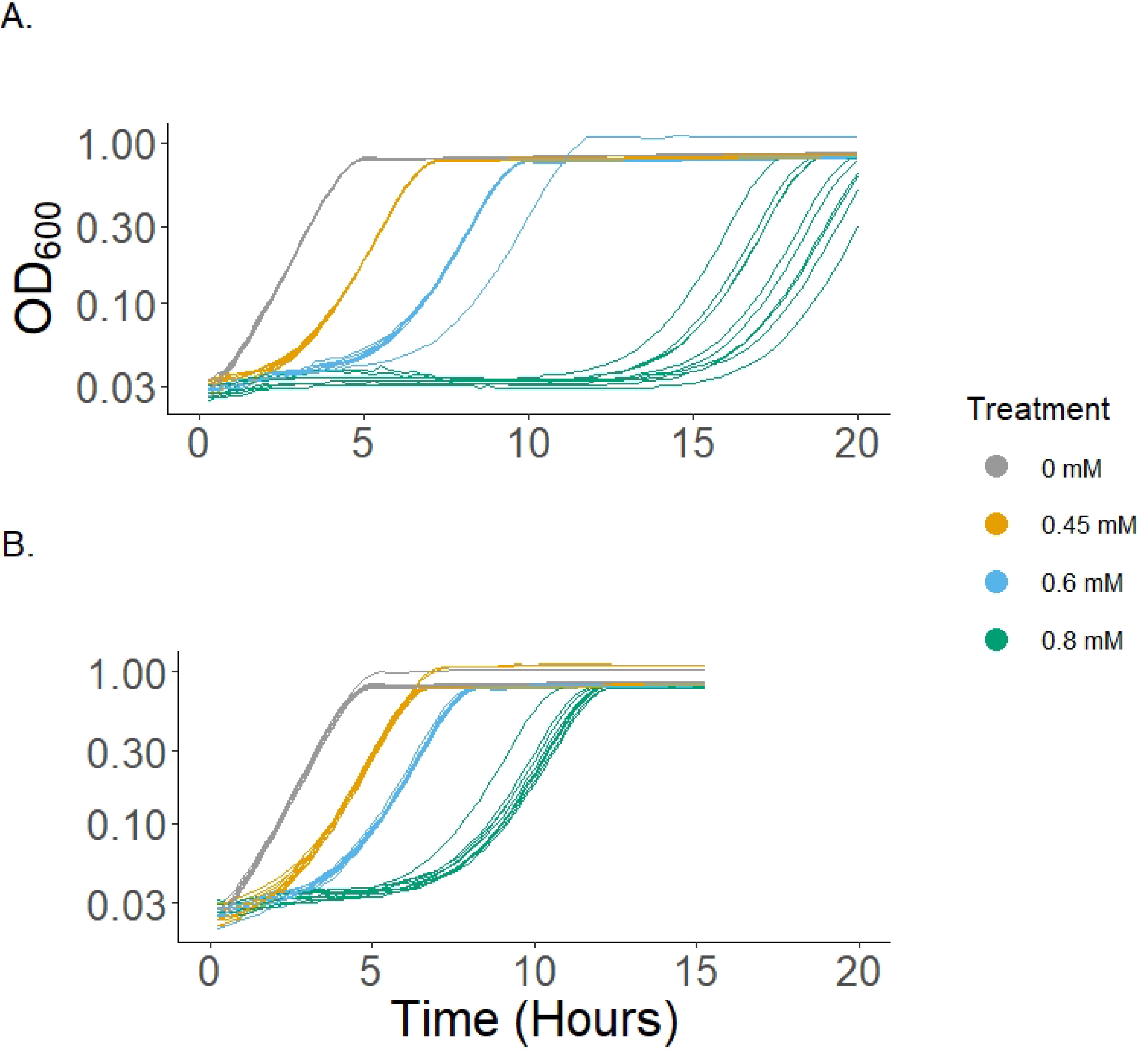
Lag time is dependent on formaldehyde concentration. As formaldehyde concentration rises, the lag time and its variability increase while growth rate is only slightly affected (S) for both (A) naïve cultures and (B) cultures pre-grown with 0.8 mM formaldehyde added. The growth curves of cultures in varying amounts of formaldehyde demonstrates the increased lag associated with higher concentration of formaldehyde, as well as increased variability between wells of the same treatment with regards to the lag (S3 Table). Pre-growth in formaldehyde prior to inoculation to some degree attenuates these dynamics (S4 Table).

### Lag times increase with increasing formaldehyde, as does their variance

Growth curves obtained by growing liquid cultures in various concentrations of formaldehyde revealed differences in lag time and its variance, but not in growth rates. Increasing the concentration of the formaldehyde in a well greatly increased the length of time that that culture spent in lag phase, as well as the variance between replicate wells in the time to reach mid-exponential growth. Cultures inoculated with cells that had been pre-grown in formaldehyde, however, did not have as great a response to increased formaldehyde in the media: they had shorter lag phases and less variance of time to mid log (S3 Table and S4 Table). Note that the observed lag times on the automated plate reader were shorter than in closed culture tubes (see Figure 3), probably due to increased oxygenation and/or formaldehyde volatility.

**Figure 3.**
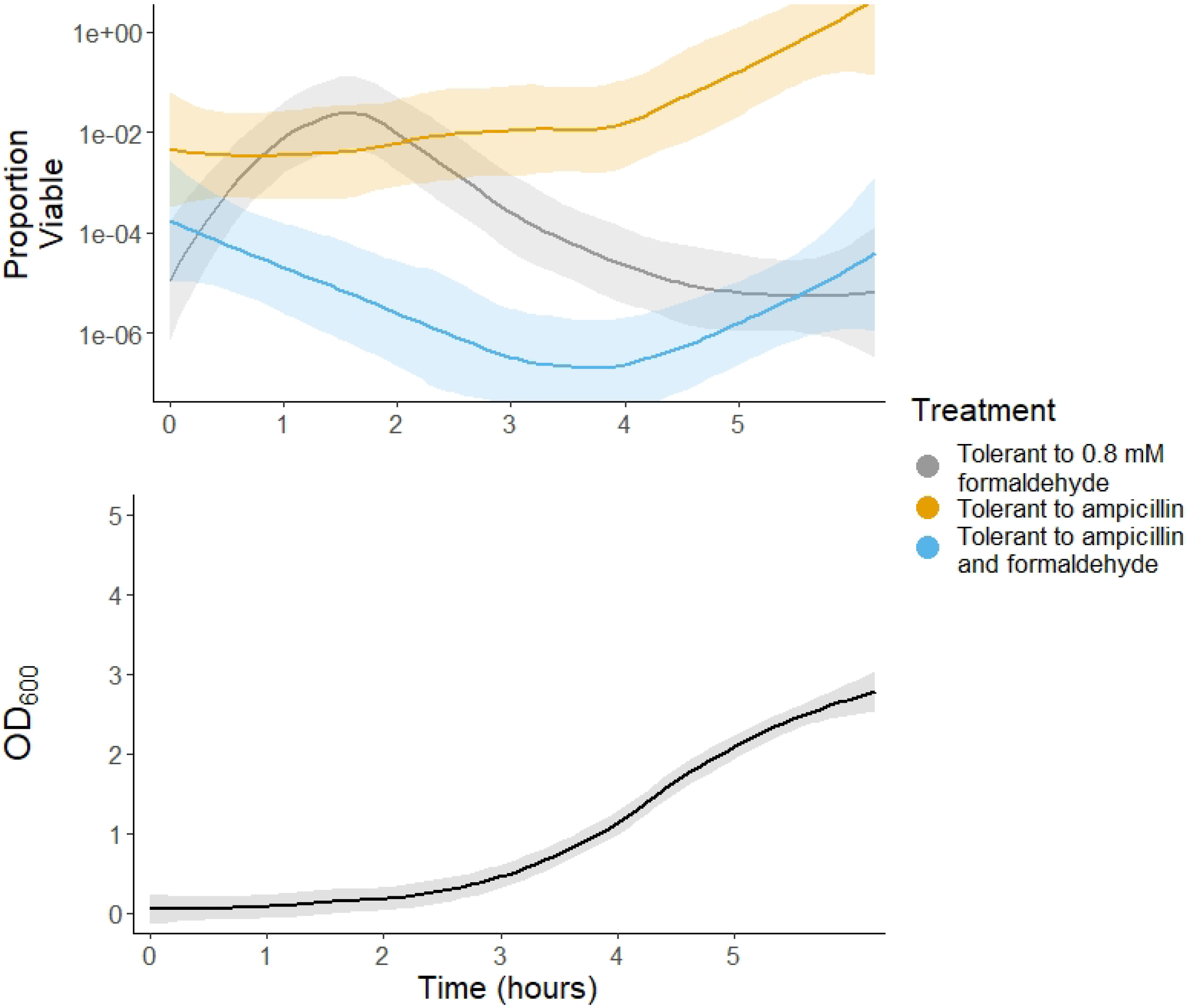
Formaldehyde is degraded before growth ensues. (A) Growth was undetectable before formaldehyde concentrations (B) decline to below about 0.1 mM, with some cultures beginning to recover before others, and a sharp decline in formaldehyde always precedes growth. (C) Plotting the phase space of OD_600_ against formaldehyde concentration shows the dependence of growth on low formaldehyde concentrations. Triangles in panel B depict the formaldehyde concentration and OD_600_ of inoculated media that did not experience growth before hour 42.

### Culture growth occurs only after substantial formaldehyde degradation

The long delay in growth of *E. coli* led us to question whether growth preceded the degradation of formaldehyde, as with *M. extorquens*. Initially the formaldehyde, as measured via colorimetric Nash assay, decreased slowly, likely as a result of glutathione-dependent oxidation of formaldehyde as discussed below, without a detectable increase in cell density. Only later in the time course did formaldehyde degradation accelerate and an increase in OD_600_ was measurable. This, along with other data (see S) suggests that formaldehyde has a bacteriostatic effect on *E. coli*, and that *E. coli* populations, unlike those of *M. extorquens*, must degrade formaldehyde to a substantial extent before growth can begin. Indeed, despite the varying times of arisal, all cultures followed remarkably similar trajectories in the OD_600_ versus formaldehyde phase space (Figure 3C). Cultures that failed to clear the media of formaldehyde, conversely, did not grow. The fact that growth only occurs after formaldehyde degrades is evocative of the dynamics of persisters in antibiotics [28], where actively growing cells are killed and non-growing cells are spared, allowing them to transition out of their non-growing state and reestablish the population following the degradation of antibiotic.

### Persistence is not responsible for formaldehyde tolerance

Because cultures were found to grow only after the majority of formaldehyde was degraded, we tested whether this quiescence indicates that formaldehyde tolerance is a pleiotropic consequence of antibiotic persistence. To investigate this, timepoints from cultures grown in media with either no or 0.8 mM formaldehyde were treated either with ampicillin for 2.5 hrs (to select for persisters, S2 Figure) or no ampicillin, and then each of these were plated to MOPS glucose medium with or without 0.8 mM formaldehyde. If persistence was the cause of formaldehyde tolerance, the dynamics of ampicillin survival should parallel formaldehyde tolerance. At t_0_ tolerance to ampicillin (Figure 4) was high due to cells having been inoculated from stationary phase, but tolerance to formaldehyde was low. By the time the cultures just began to show net growth via OD_600_, ampicillin survival had not yet dropped but formaldehyde tolerance greatly increased. As growth began, ampicillin tolerance decreased, formaldehyde tolerance remained high, and nearly all ampicillin-tolerant cells could also grow on formaldehyde. The initial lack of ampicillin tolerance in media lacking formaldehyde is likely due to growth commencing at inoculation; for the formaldehyde exposed cultures this effect is suppressed by the presence of formaldehyde in the media (see Figure 3). During midexponential phase, formaldehyde tolerance then began to drop. As the cultures entered stationary phase again, ampicillin survival increased and converged to 100% of the population as the culture approached stationary phase. By this point in time, the overwhelming majority of formaldehyde tolerant individuals were also non-growing cells.

**Figure 4.**
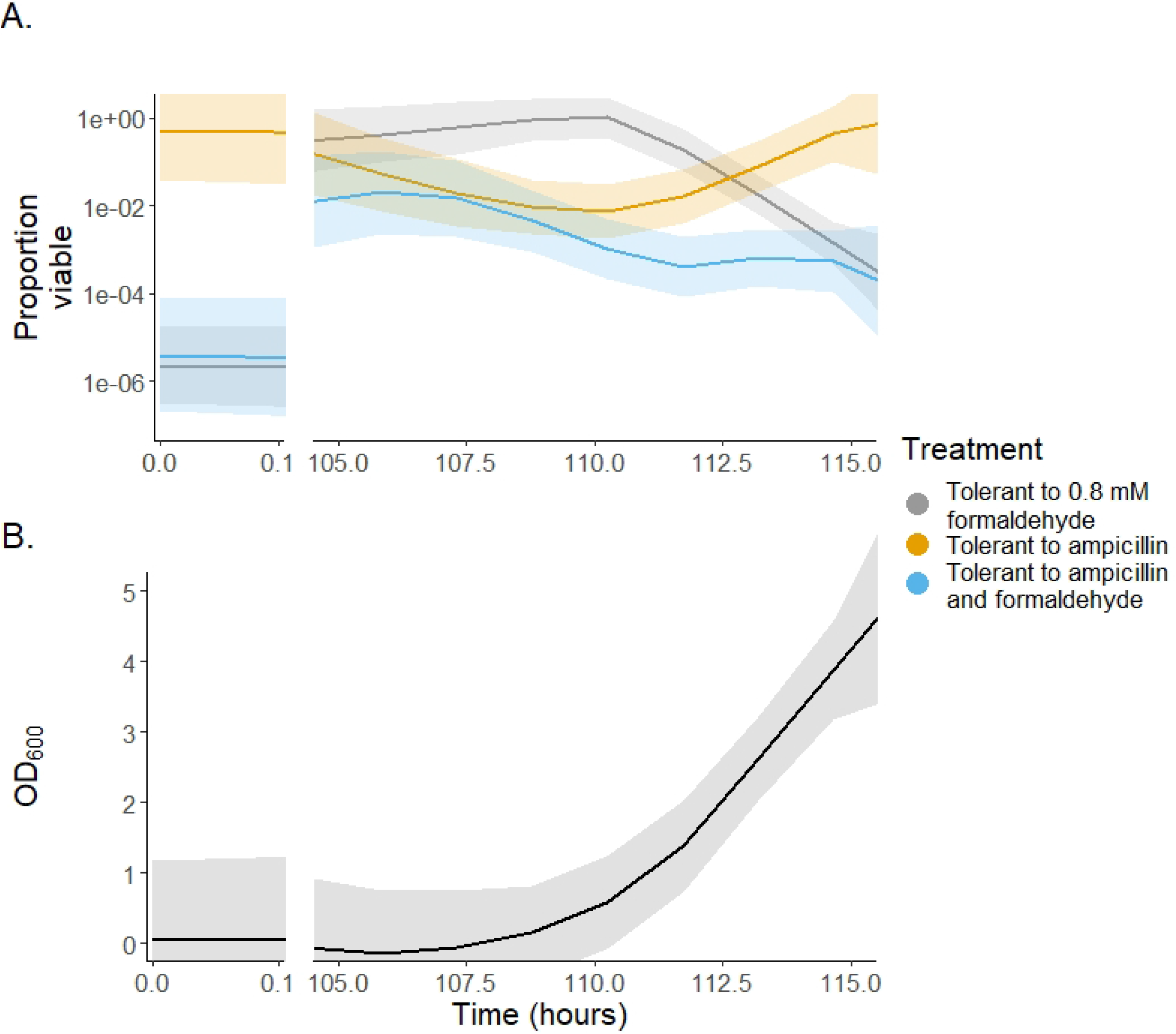
Persistence does not explain formaldehyde tolerance. Media with formaldehyde was inoculated, and at the commencement of growth samples from the cultures were either treated with ampicillin to select for persisters or not treated, and then each of these were periodically spot plated onto either minimal glucose medium with or without formaldehyde.. Shaded regions represent the standard error.

### Cells pre-grown in media containing formaldehyde oxidize formaldehyde at a greater rate

Because of the observations of reduced lag in formaldehyde for cultures pre-grown in media with formaldehyde (Figure 2) and of growth occurring only following formaldehyde degradation (Figure 3), we hypothesized that exposure to formaldehyde induces a phenotype that degrades formaldehyde faster. To test this, we measured the formaldehyde-dependent rate of formation of NADH by whole cell lysates and found that the cultures grown in formaldehyde did oxidize formaldehyde at a much higher rate (Fig. 5). The observed apparent induction of formaldehyde oxidizing genes is consistent with previous studies that observed increases in induction of the *frmAB* operon, which contains genes responsible for glutathione-dependent oxidation of formaldehyde [29].

**Figure 5.**
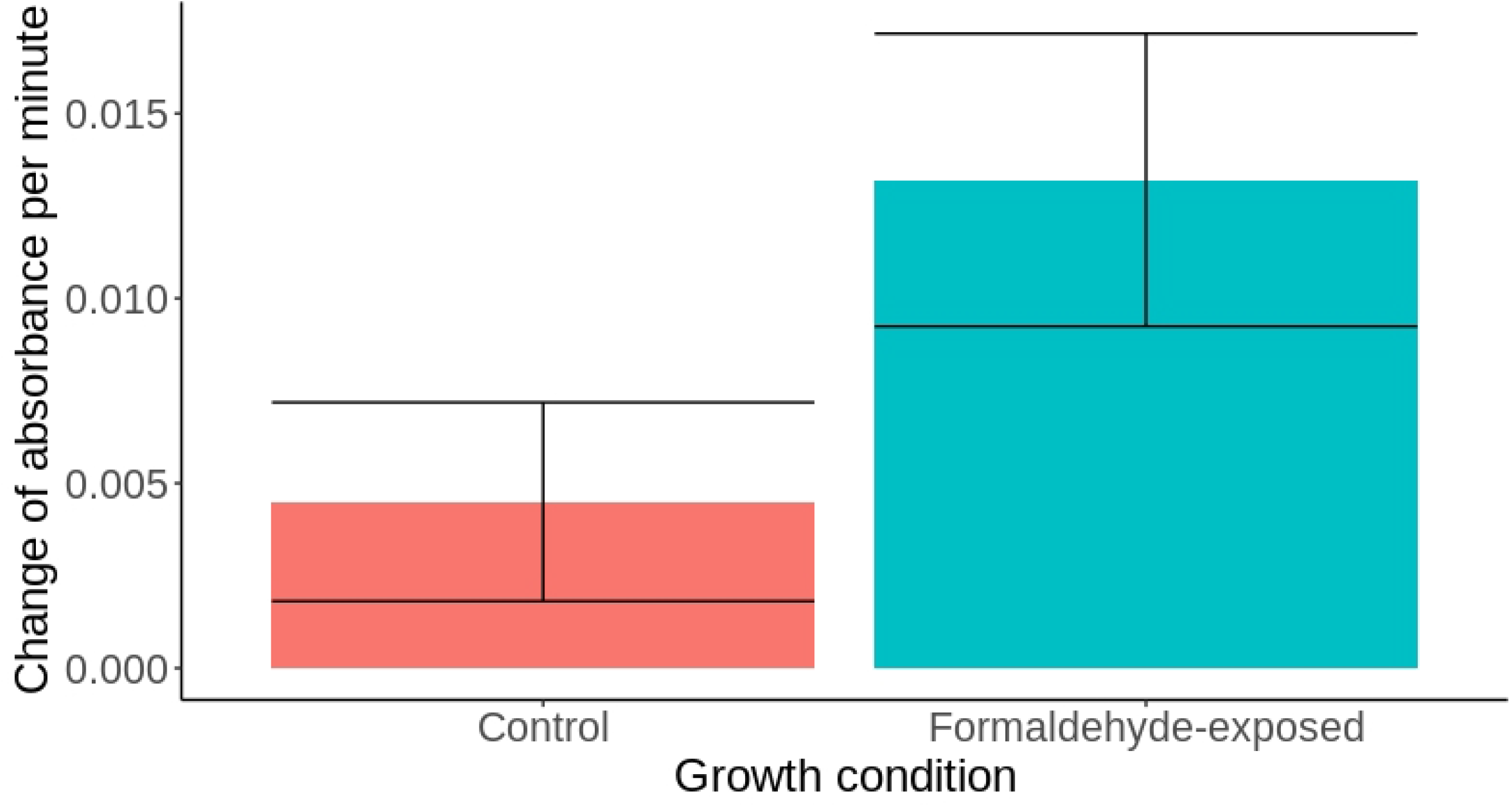
Formaldehyde oxidation is more rapid in cells previously exposed to formaldehyde. The formaldehyde oxidation rate of cultures pre-grown in formaldehyde and another not previously exposed to formaldehyde were measured via colorimetric assay of NADH production. The rate of formaldehyde oxidation is much greater in cells grown in the presence of formaldehyde.

### Formaldehyde-grown cultures retain formaldehyde tolerance longer than naive cultures

To test the idea that dormant cells can preserve the formaldehyde tolerant phenotype, and that the increase in formaldehyde tolerance in formaldehyde naive cells (Figure 4) is due to active growth, we sampled cultures at about four days following their entrance into stationary phase and spot plated them onto plates containing 0.8 mM formaldehyde. As seen in Figure 6, after four days the formaldehyde naive culture had relatively little viability compared to the formaldehyde exposed culture. This is congruent with our other results showing that cultures grown in the absence of formaldehyde have low tolerance while in stationary phase. It also indicates that a formaldehyde tolerant cell, once dormant, will retain this tolerance for a considerable amount of time.

**Figure 6.**
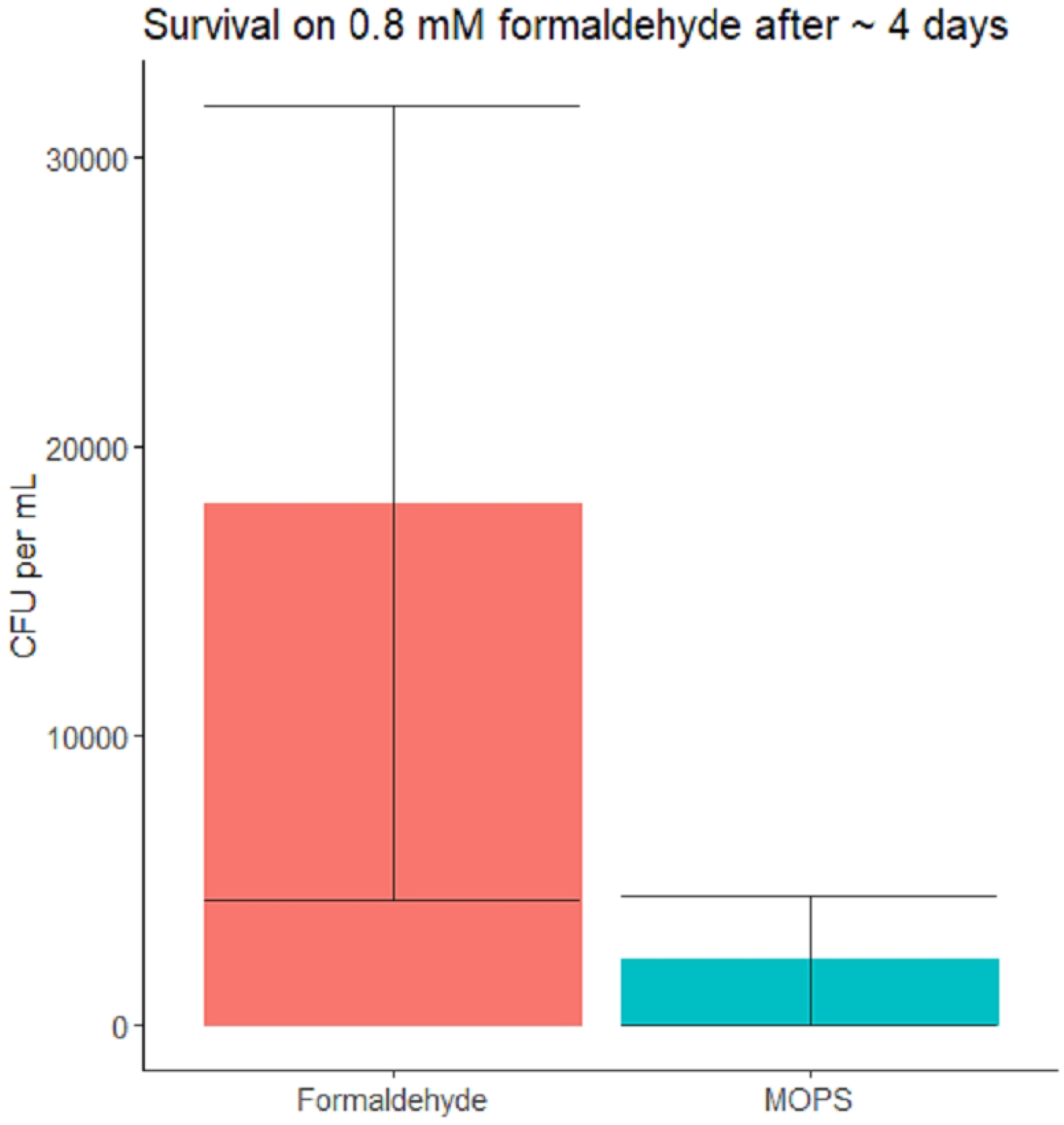
Tolerance to formaldehyde of two cultures after 4 days in stationary phase. Cultures grown in media containing formaldehyde maintain greater tolerance to formaldehyde after four days than cultures that were not. Cells which had been grown in media containing formaldehyde showed noticeably greater tolerance to formaldehyde even after four days in stationary.

**Figure 7.**
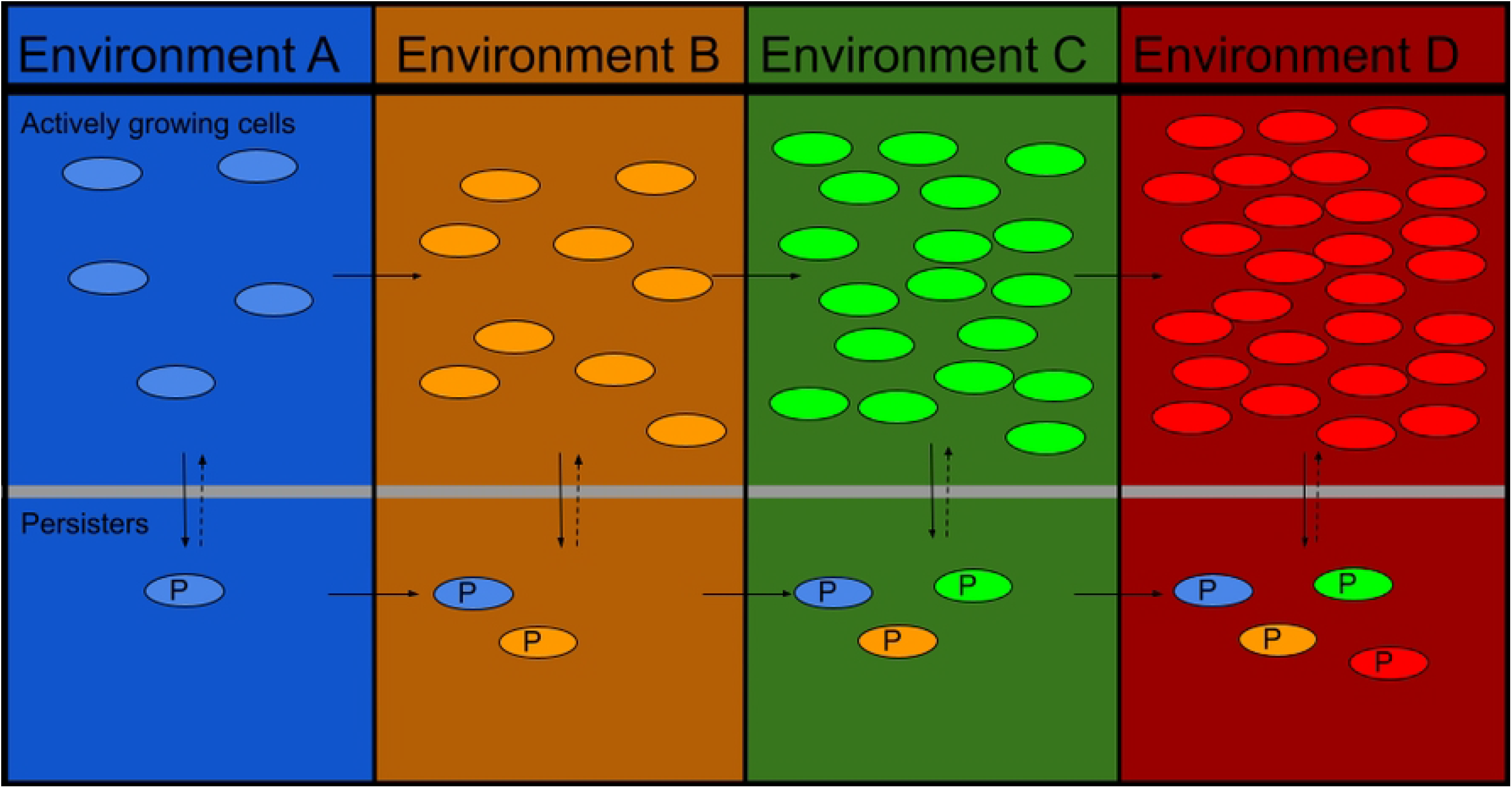
A schematic of the proposed mechanism by which persistence generates metabolic heterogeneity. As the environment changes, actively growing cells express new phenotypes, as indicated by the colored backgrounds and cells. Some growing cells however produce persisters (lower row, cells marked with a P), which are much less metabolically active and do not change their phenotype. This allows a population to retain phenotypic memory longer than would be possible through typical phenotypic inheritance.

## Discussion

Here we have uncovered that the non-methylotrophic model organism *E. coli*, like the methylotroph *M. extorquens*, exhibits substantial phenotypic heterogeneity in formaldehyde tolerance. Although external formaldehyde may not be a frequent stressor for *E. coli*, it impinges upon cell physiology in several ways, notably in generating protein damage [30]. Given that the formaldehyde tolerance first observed in *M. extorquens* is correlated with, and perhaps caused by, upregulation of conserved proteins that are responsible for dealing with general oxidative stress and protein damage, it seemed plausible that non-methylotrophs might also exhibit phenotypic heterogeneity despite not facing formaldehyde stress as a part of their central metabolism. Although the work presented here confirms this to be the case, with multiple features of heterogeneity in formaldehyde tolerance being similar between the two organisms, there are also myriad differences in the manifestation of formaldehyde stress tolerance in *E. coli*.

### Continuous, dynamic heterogeneity in tolerance

The most fundamental similarity between the formaldehyde tolerance phenomena in *E. coli* and *M. extorquens* was that both organisms display a wide, smooth continuum in the formaldehyde concentrations that individual cells can manage to grow in. In both organisms, small increases in formaldehyde concentration have undetectable effects on cell viability. If formaldehyde concentrations are increased further, populations of either organism exhibit an effectively exponential decline in cell viability. Although rare (10^-7^), some cells survived and grew at 0.8 mM formaldehyde, which was twice as high as the concentration which first generated a significant drop in viability, 0.4 mM. A similar proportional range of tolerance was found for *M. extorquens*, whereby the highest concentration naïve cells were found to tolerate (5 mM) was 2.5-fold as high as the first concentration that was lethal for some of the population [17]. Thus, despite a nearly order of magnitude greater tolerance in the methylotrophic organism with high levels of cytoplasmic formaldehyde oxidation capacity [17], the non-methylotrophic *E. coli* displayed a similarly wide, continuous range of heterogeneity in tolerance.

A further parallel between the two species was that, following growth in media containing formaldehyde the distributions of tolerance shifted upwards towards increased tolerance to formaldehyde. This shows that the observed phenotype is inducible; both bacteria are capable of upregulating a response to deal with formaldehyde stress. The proportion of *E. coli* cells that could tolerate growth at 0.6 or 0.8 mM formaldehyde rose two to three orders of magnitude upon formaldehyde exposure. As with *M. extorquens*, however, the tolerance distribution of the population rapidly returned to its naïve distribution upon a cycle of regrowth in minimal glucose medium in the absence of formaldehyde. This suggests that, as was demonstrated for *M. extorquens*, the highly tolerant *E. coli* cells were not genetic variants, but simply exhibited a change in phenotype.

A subtle, second parallel between the two organisms was that exponential phase cells had a higher tolerance distribution than stationary phase cells. There was enhanced tolerance to formaldehyde in *E. coli* cultures that had just begun growing even in the absence of external formaldehyde (see S6 Figure). It is possible that during rapid growth, deformylation of *N*-formylmethionine can cause transient spikes of formaldehyde intracellularly, inducing greater tolerance to formaldehyde than stationary cultures, or that this increase in tolerance is actually due to more general mechanisms, for instance upregulation of chaperonins during the transition from stationary to growing states. Regardless, the increased tolerance of growing cells compared to stationary phase cells runs counter to the general tendency for non-growing cells to have greater resistance to a variety of stressors.

### *E. coli* cultures require the degradation of formaldehyde before growth can commence, unlike *M. extorquens*

While *M. extorquens* cultures treated with higher and higher formaldehyde levels took longer for growth to become observable, replicate cultures remained remarkably similar in their timing [17]. As with *M. extorquens*, sub-lethal concentrations of formaldehyde were found to generate an extended lag for *E. coli*, rather than to lead to a slower rate of growth. We found that *E. coli* cultures exposed to higher formaldehyde concentrations displayed substantial differences in the timing for which growth would ensue. At the 0.8 mM exposure commonly used, some cultures never recovered at all (Figure 3).

These variable dynamics appear to relate to the fact that *E. coli* growth only becomes detectable after the formaldehyde concentrations are significantly reduced. Whereas the viable count of *M. extorquens* growth (after a period of death) was found to begin well before formaldehyde had begun to decline, here we found that the variable timing of optical density increase in *E. coli* cultures seemed to correlate with the timing of when populations managed to detoxify their medium. It should be noted that there was a clear acceleration of formaldehyde degradation immediately prior to growth in the *E. coli* cultures. This may be due to a small increase in culture density below detection, upregulation of formaldehyde detoxification pathways, or an increased physiological capacity to oxidize formaldehyde as the concentrations become lower.

The inability of *E. coli* to initiate growth until formaldehyde is quite low, unlike *M. extorquens*, may be due to the fact that the latter possesses the recently-discovered formaldehyde sensor, EfgA, which acts via decreasing protein translation in the cell when it binds formaldehyde [31]. EfgA aids *M. extorquens* in managing the transition from non-methylotrophic to methylotrophic growth, a period of time when formaldehyde transiently accumulates [32]. *M. extorquens* lacking *efgA* have an increased lag during this transition due to heterogeneity in the ability of single cells to initiate growth. Conversely, introduction of EfgA into *E. coli* was found to decrease the lag time of cultures exposed to formaldehyde and raise their tolerance level [33]. In the absence of EfgA to modulate translation, either directly or indirectly, *E. coli* cultures may be generating too many targets for formaldehyde damage early in the transition to growth in its presence.

### Prior exposure to formaldehyde induces a greater rate of formaldehyde oxidation in whole-cell extracts

Assaying enzyme activity in cultures näive to and grown in the presence of formaldehyde suggested induction of enhanced formaldehyde oxidation (Figure 5). This apparent plasticity in formaldehyde oxidation, combined with the formaldehyde concentration dependence of growth (Figure 3) is evidence that the enhanced growth of cultures pre-exposed to formaldehyde (Figure 2) is the result of increased formaldehyde oxidation. Furthermore, the heterogeneity of formaldehyde tolerance (Figure 1) may be the result of differences in the capacity of individuals to oxidize formaldehyde. Such behavior has been observed in the context of chloramphenicol tolerance [34], where individuals resistant to chloramphenicol exhibited bistable growth near the minimum inhibitory concentration of chloramphenicol. Individuals with sufficient levels of a chloramphenicol detoxifying enzyme were capable of establishing colonies, while individuals that had insufficient concentrations were trapped in a non-growing state.

### Phenotypic heterogeneity in formaldehyde tolerance is unrelated to persistence

The fact that growth only took place after a substantial reduction in formaldehyde concentrations raised the possibility that the survivors that represent formaldehyde tolerance were simply persisters that recovered growth after the stressor was sufficiently removed by non-growing but metabolically active cells. By simultaneously tracking persistence to ampicillin with formaldehyde tolerance we were able to rule this out. Persistence levels were high prior to growth initiating, dropped during growth, and then rose again, as expected. In contrast, formaldehyde tolerance rose from exceedingly rare to high when growth initiated, remained high for a period of growth, and then began to drop during exponential phase with no recovery as the culture re-entered stationary phase. The uncorrelated, generally opposite dynamics of formaldehyde tolerance and persistence clearly indicate that they represent independent phenotypic states. The presence of cells who could survive ampicillin exposure and recover growth on formaldehyde demonstrated that they are not mutually exclusive phenotypic states.

### Persistence can “preserve” the acquired formaldehyde tolerance phenotype

Although formaldehyde tolerance is not a byproduct of persistence, the latter phenotypic state appears to be capable of locking in and preserving heterogeneity in the former. During the period that the majority of the population had high formaldehyde tolerance (see Figure 6B), the persisters generated, while rare in the population, were also formaldehyde tolerant. However, as the culture entered stationary phase, the only formaldehyde tolerant cells left were also simultaneously persisters. This is likely caused by the cells in a formaldehyde tolerant state transitioning directly into a persistent state. Retesting the culture ~4 days later (Figure 6) revealed that the formaldehyde tolerance phenotype was maintained through this time, indicating that dormant cells can retain phenotypic memory of the state they were in when they entered dormancy, whether that dormancy is induced by persistence or starvation.

Discovering that persistence can preserve phenotypic formaldehyde tolerance suggests that persisters may be able to preserve cells acclimated to a wide variety of recently encountered stressors. To our knowledge, the concept that persisters can preserve heterogeneity in other phenotypes in a population has not been suggested previously. Whereas cultures that continue to grow lose enhanced formaldehyde tolerance fairly rapidly, likely due to dilution by growth of proteins involved, persisters can maintain this heterogeneity, likely through locking into place differences in protein expression. In environments that fluctuate between times of growth and stress, this may be a critical source of variation – at the phenotypic level – that allows cells to survive a wider variety of stresses than realized.

## Supplementary Information

**S1 Table. Formula for MOPS minimal media.**

This buffer is 10x strength, and is stored at 4 °C separate from its phosphate and carbon source until needed. The pH is brought to 7.4 with NaOH, and filter sterilized. Phosphate is supplied in the form of KH_2_PO_4_ at a concentration of 1.32 mM, and glucose is supplied at a concentration of 11.1 mM.

**S2 Figure. Killing curve for ampicillin.**

Samples from an *E. coli* culture at mid-exponential phase were subjected to treatment with 100 μg/mL ampicillin at 37 °C for various amounts of time. Following treatment, the cells were spun down, their supernatant poured off, and resuspended in an equivalent volume. They were then spot plated on permissive plates.

**S3 Table. Mean time and standard deviation to reach an OD_600_ of 0.4 for a formaldehyde naive culture.**

For each growth curve, the time point before the culture reached an OD_600_ equal to or greater than 0.4 was averaged and the standard deviation calculated for each treatment. Timepoints occured in 15 minute increments. This table corresponds to the growth curves in Figure 2 A.

**S4 Table. Mean time and standard deviation to reach an OD_600_ of 0.4 for a formaldehyde pre-exposed culture.**

For each growth curve, the time point before the culture reached an OD_600_ equal to or greater than 0.4 was averaged and the standard deviation calculated for each treatment. Timepoints occured in 15 minute increments. This table corresponds to the growth curves in Figure 2 B.

**S5 Figure. Growth rates of cultures grown with or without formaldehyde present in media.**

The growth rates of cultures naive to formaldehyde stress (panel A.) or grown in the presence of 0.8 mM formaldehyde (panel B) after inoculation into media containing various concentrations of formaldehyde. This illustrates that although the differences in growth rate are statistically significant (as assessed by Tukey’s HSD test, p < 0.05) they are relatively small.

**S6 Figure. Proportion of naive cells tolerant to ampicillin, formaldehyde or both**.

Unlike in figure 4, in which the cells were grown in media with formaldehyde, the growth media for this culture did not contain formaldehyde. Unlike in Figure 4, the population which is both formaldehyde and ampicillin tolerant is far smaller than either population which is tolerant to each challenge by itself, except at inoculation or entrance to stationary phase.

**S7 Figure. CFU of ampicillin/formaldehyde tolerance experiment.**

Rather than the proportion of cells tolerant to formaldehyde, ampicillin or both at each timepoint as in Figure 4, this graph displays the actual CFU of all four treatments. Panel A depicts the growth of the populations in media that did not contain formaldehyde, while panel B depicts growth in media that did contain formaldehyde, and the shaded areas represent the standard error.

## Notes

### Competing Interest Statement

The authors have declared no competing interest.

